## Supplementary Tables and Figures for "Brain-derived estrogens facilitate male-typical behaviors by potentiating androgen receptor signaling in medaka"

**Supplementary file 1. Abbreviations of brain nuclei.**

| abbreviation | full name | location |
|---|---|---|
| aNVT | anterior part of NVT | hypothalamus |
| aPMp | anterior part of PMp | preoptic area |
| aPPp | anterior part of PPp | preoptic area |
| NAT | anterior tuberal nucleus | hypothalamus |
| NPT | posterior tuberal nucleus | hypothalamus |
| NRL | lateral recess nucleus | hypothalamus |
| NVT | ventral tuberal nucleus | hypothalamus |
| PMg | gigantocellular portion of the magnocellular preoptic nucleus | preoptic area |
| PMm | magnocellular portion of the magnocellular preoptic nucleus | preoptic area |
| PMp | parvocellular portion of the magnocellular preoptic nucleus | preoptic area |
| pNVT | posterior part of NVT | hypothalamus |
| PPa | anterior parvocellular preoptic nucleus | preoptic area |
| pPMp | posterior part of PMp | preoptic area |
| PPp | posterior parvocellular preoptic nucleus | preoptic area |
| pPPp | posterior part of PPp | preoptic area |
| SC | suprachiasmatic nucleus | preoptic area |
| VM | ventromedial nucleus | thalamus |
| Vp | posterior nucleus of the ventral telencephalic area | ventral telencephalon |
| Vs | supracommissural nucleus of the ventral telencephalic area | ventral telencephalon |
| Vv | ventral nucleus of the ventral telencephalic area | ventral telencephalon |

**Supplementary file 2. Primers and probes used in this study.**

| primer/ probe | target | direction | purpose | sequence (5′ to 3′) |
|---|---|---|---|---|
| primer | *cyp19a1b* | forward | genotyping (gDNA PCR) | GACTTGGTCCTGTCCTGTCCTA |
| primer | *cyp19a1b* | reverse | genotyping (gDNA PCR) | ATCCTGGTTTTCTTCCACAGAG |
| primer | *cyp19a1b* | forward | genotyping (CS) | TTGTGAGGGTATGGATTAATGG |
| primer | *cyp19a1b* | forward | genotyping (HRM) | CGGCTGAAAGCTTGTTTACCTA |
| primer | *cyp19a1b* | reverse | genotyping (HRM) | CCCGAATCTAGACGTGTAGTGG |
| probe | *cyp19a1b* | forward | genotyping (HRM) | TGAGGTGTACCATGTTTTGAAGAGC |
| primer | *cyp19a1b* | forward | real-time PCR | AAGAAGATGATCCAGCAAGAG |
| primer | *cyp19a1b* | reverse | real-time PCR | AGCATCAGAAGAAGTAAGAAAAGTG |
| primer | *esr2a* | forward | genotyping (gDNA PCR) | ATGTCGCTTTTGCAGTTTAAGCTG |
| primer | *esr2a* | reverse | genotyping (gDNA PCR) | ATGAACACGGATCTGCTGATGG |
| primer | *esr2a* | forward | genotyping (CS) | ACGGCTTTGAAGATCCTTGGCT |
| primer | *esr2a* | forward | genotyping (HRM) | CAGGCGGCAAGTCTGAACTC |
| primer | *esr2a* | reverse | genotyping (HRM) | CTCCATTTTACCTTGGATGCTCC |
| probe | *esr2a* | forward | genotyping (HRM) | ATACCACTACGGCGTGTGGTCATGCGA G |

gDNA PCR, PCR on genomic DNA; CS, cycle sequence; HRM, high-resolution melting analysis.

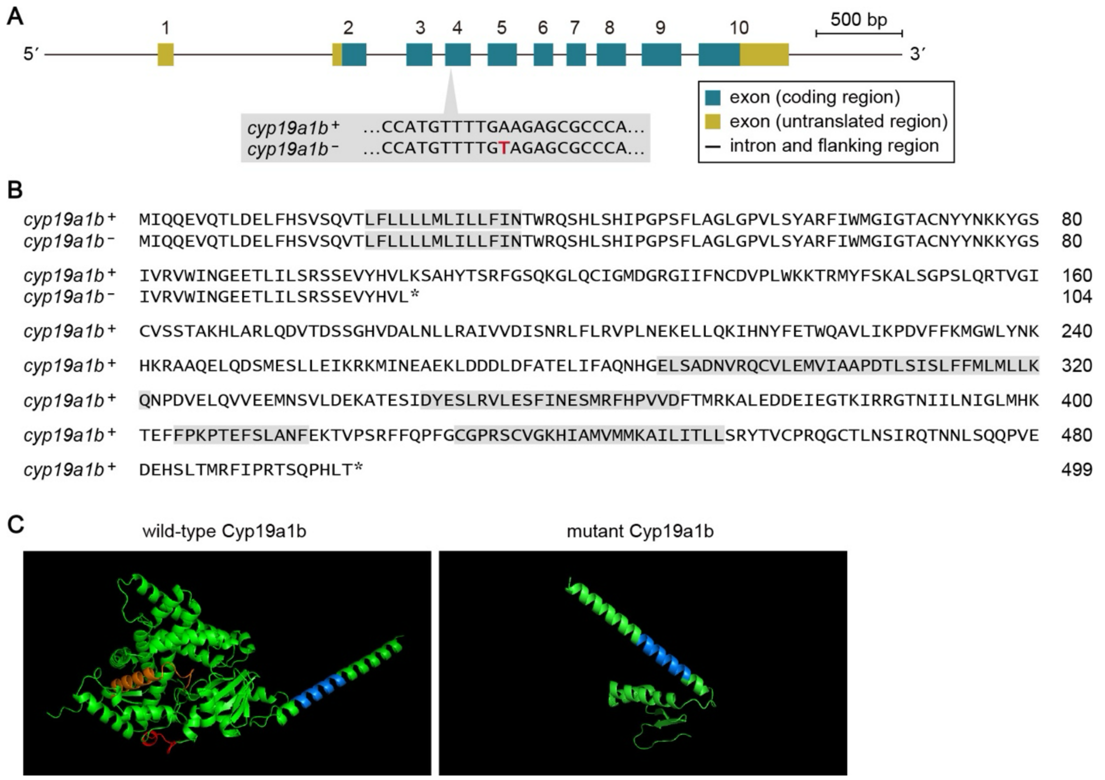


**Figure 1—figure supplement 1. Generation of *cyp19a1b*-deficient medaka.**

(**A**) Schematic of the *cyp19a1b* locus showing the location of the mutation identified by TILLING. Each exon is numbered. Nucleotide sequences of the wild-type (*cyp19a1b*$^{+}$) and mutant (*cyp19a1b*$^{-}$) alleles at the mutation site are shown, with the substituted nucleotide in red. (**B**) Comparison of the amino acid sequences deduced from the *cyp19a1b*$^{+}$ and *cyp19a1b*$^{-}$ alleles. Amino acid numbers are shown on the right. Putative functional domains (membrane-spanning region [residues 21–34 of wild-type Cyp19a1b], I-helix region [287–321], Ozol's peptide region [346–368], aromatic region [404–415], and heme-binding region [429–452]) are shaded in gray. Asterisks denote stop codons. (**C**) Predicted three-dimensional structures of wild-type (left) and mutant (right) Cyp19a1b proteins. Key structural features are annotated as follows: membrane helix (blue), aromatic region (red), and heme-binding loop (orange).

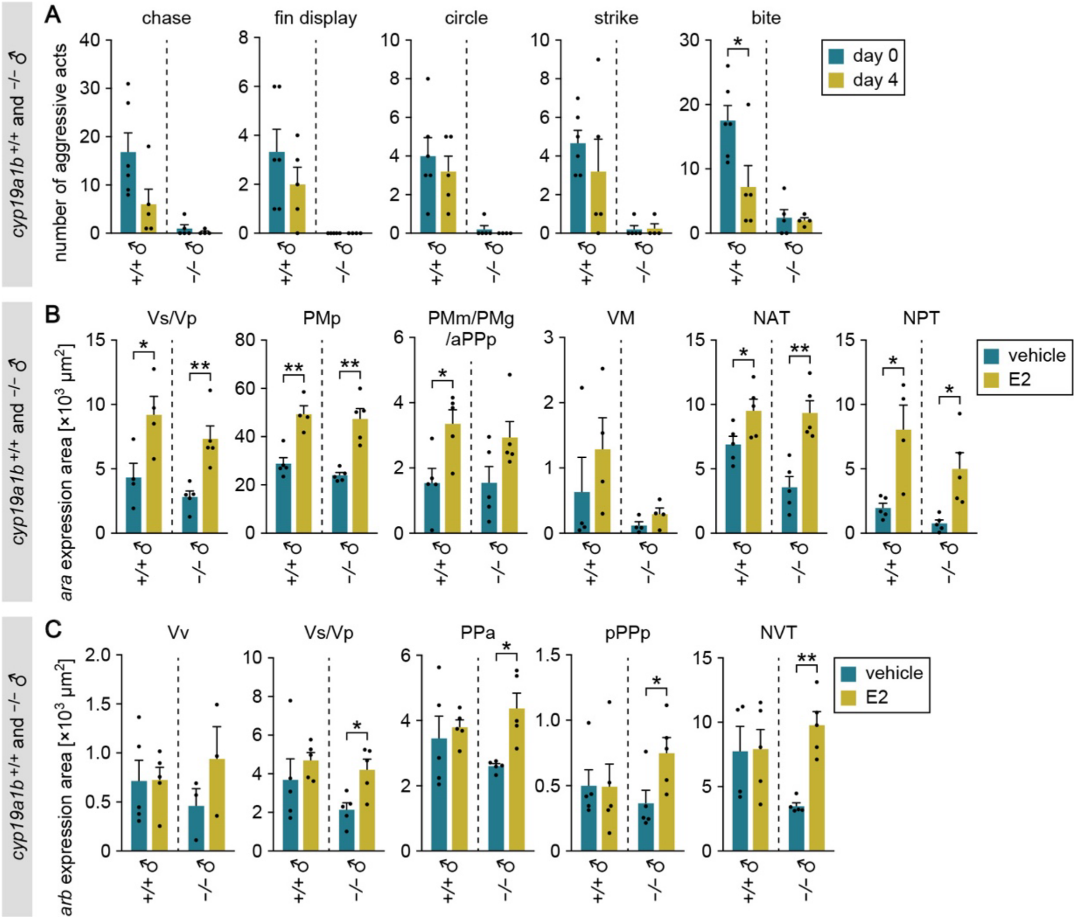


**Figure 2—figure supplement 1. Effect of estrogen replacement on aggression and brain *ara* and *arb* expression in *cyp19a1b*-deficient males.**

(**A**) Total number of each aggressive act observed among *cyp19a1b*$^{+/+}$ or *cyp19a1b*$^{-/-}$ males before (day 0; n = 6 and 5, respectively) or after (day 4; n = 5 and 4, respectively) E2 treatment. (**B**) Total area of *ara* expression signal in the Vs/Vp, PMp, PMm/PMg/aPPp, VM, NAT, and NPT of *cyp19a1b*$^{+/+}$ and *cyp19a1b*$^{-/-}$ males treated with vehicle alone or E2 (n = 5 per group except for Vs/Vp of vehicle- and E2-treated *cyp19a1b*$^{+/+}$ males, PMp and NPT of E2-treated *cyp19a1b*$^{+/+}$ males, and VM of all males, where n = 4). (**C**) Total area of *arb* expression signal in the Vv, Vs/Vp, PPa, pPPp, and NVT of *cyp19a1b*$^{+/+}$ and *cyp19a1b*$^{-/-}$ males treated with vehicle alone or E2 (n = 5 per group except for NVT of vehicle-treated *cyp19a1b*$^{+/+}$ males, where n = 4; and Vv of vehicle- and E2-treated *cyp19a1b*$^{-/-}$ males, where n = 3). For abbreviations of brain nuclei, see Supplementary file 1. Statistical differences were assessed by unpaired *t* test, with Welch's correction where appropriate (A, B, C). Error bars represent SEM. $*P < 0.05$, $**P < 0.01$.

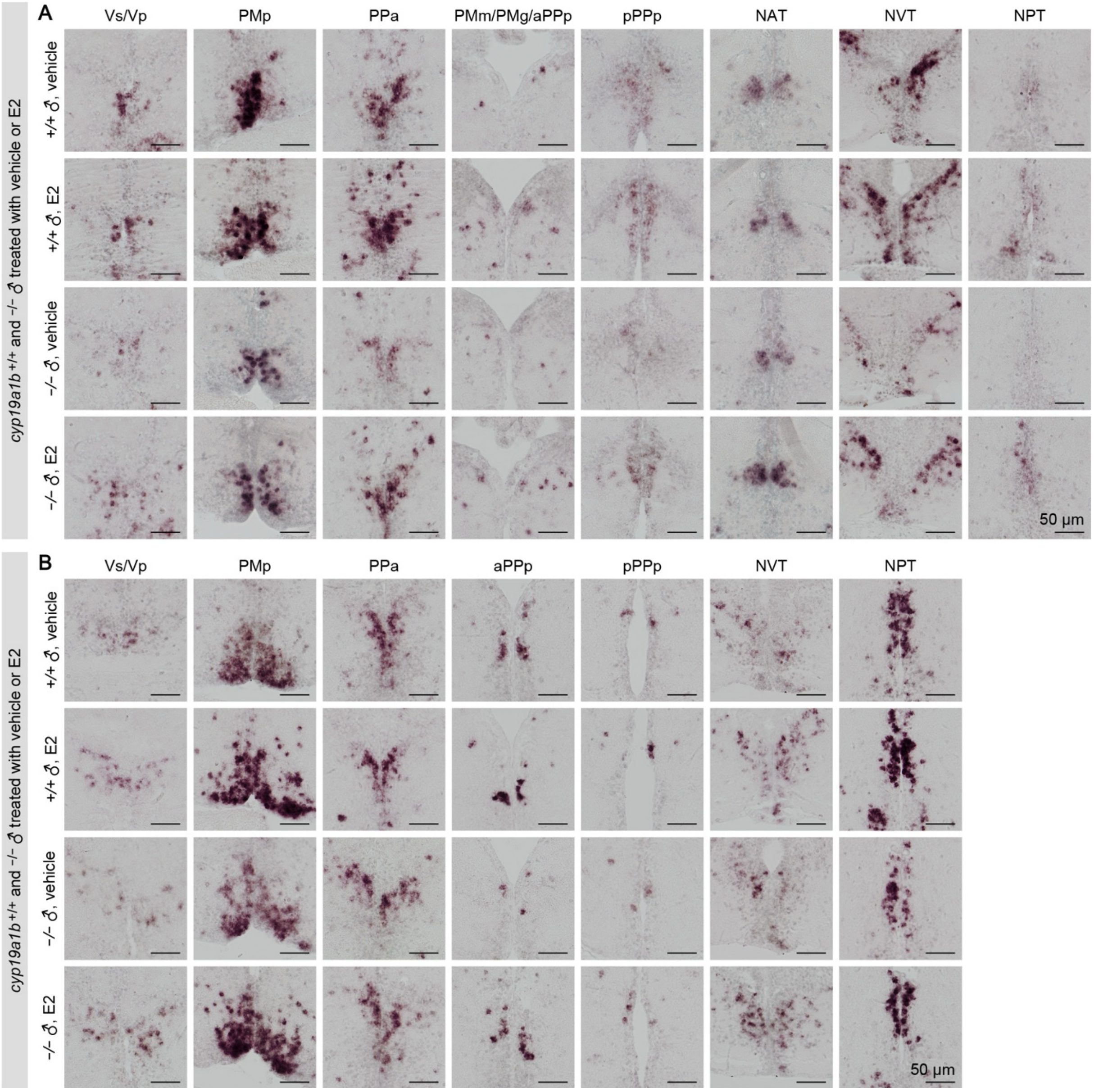


**Figure 2—figure supplement 2. Effect of estrogen replacement on brain *ara* and *arb* expression in *cyp19a1b*-deficient males.**

(**A**) Representative images of *ara* expression in the Vs/Vp, PMp, PPa, PMm/PMg/aPPp, pPPp, NAT, NVT, and NPT of *cyp19a1b*$^{+/+}$ and *cyp19a1b*$^{-/-}$ males treated with vehicle alone or E2. (**B**) Representative images of *arb* expression in the Vs/Vp, PMp, PPa, aPPp, pPPp, NVT, and NPT of *cyp19a1b*$^{+/+}$ and *cyp19a1b*$^{-/-}$ males treated with vehicle alone or E2. Scale bars represent 50 μm. For abbreviations of brain nuclei, see Supplementary file 1.

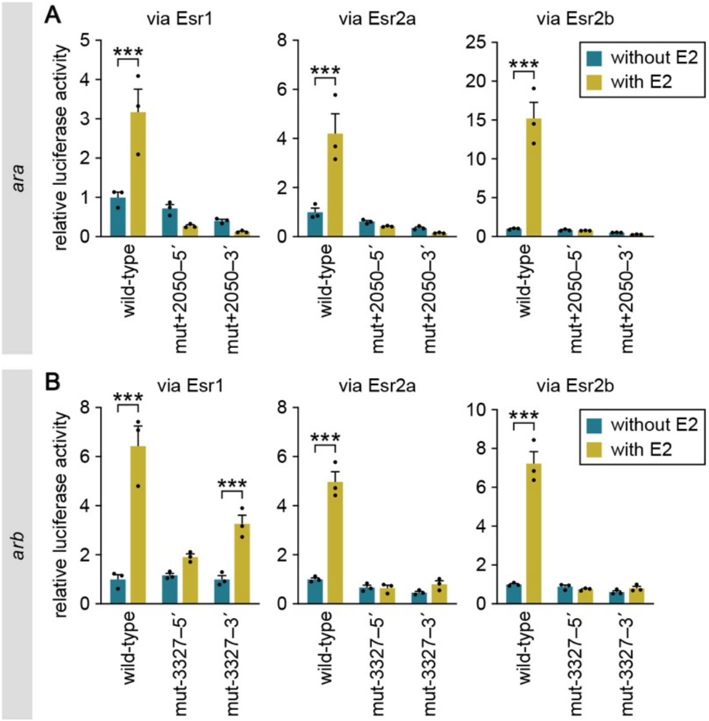


**Figure 4—figure supplement 1. Effect of mutations in each half-site of the identified ERE on the E2-induced activation of *ara* and *arb* transcription.**

(**A**) Effect of mutations in each half-site of the ERE at position +2050 of the *ara* locus on the E2-induced activation of *ara* transcription. Cultured cells were transfected with a wild-type luciferase reporter construct or a construct carrying a mutation in the 5′ (mut+2050–5′) or 3′ (mut+2050–3′) half-site of the ERE, together with an Esr1, Esr2a, or Esr2b expression construct. The cells were stimulated with or without E2, and luciferase activity was measured. (**B**) Effect of mutations in each half-site of the ERE at position –3327 of the *arb* locus on the E2-induced activation of *arb* transcription. Cultured cells were transfected with a wild-type luciferase construct or a construct carrying a mutation in the 5′ (mut-3327–5′) or 3′ (mut-3327–3′) half-site of the ERE, together with an Esr1, Esr2a, or Esr2b expression construct. The cells were stimulated with or without E2, and luciferase activity was measured. Values are represented relative to the wild-type construct without E2 stimulation. Statistical differences were assessed by unpaired $t$ test with Bonferroni–Dunn correction (A, B). Error bars represent SEM. ***$P < 0.001$.

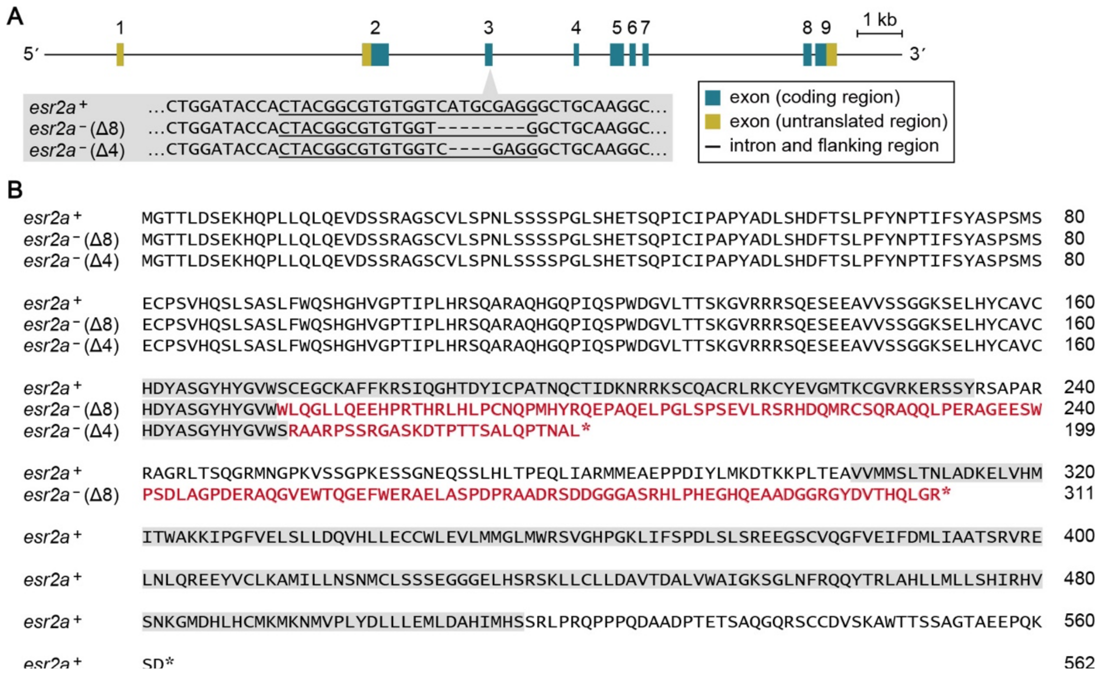


**Figure 5—figure supplement 1. Generation of *esr2a*-deficient medaka.**

(**A**) Schematic of the *esr2a* locus showing the location of the CRISPR target site. Each exon is numbered. Nucleotide sequences of the wild-type (*esr2a*$^{+}$) and mutant (*esr2a*$^{-}$ [Δ8] and *esr2a*$^{-}$ [Δ4]) alleles at the target site are shown, with the CRISPR RNA target sequence underlined. Dashes indicate deleted nucleotides. (**B**) Comparison of the amino acid sequences deduced from the *esr2a*$^{+}$ and *esr2a*$^{-}$ (Δ8 and Δ4) alleles. Amino acid numbers are shown on the right. The DNA-binding domain (residues 161–234 of wild-type Esr2a) and ligand-binding domain (304–514) are shaded in gray. The sequences altered by the frameshift are indicated in red. Asterisks denote stop codons.

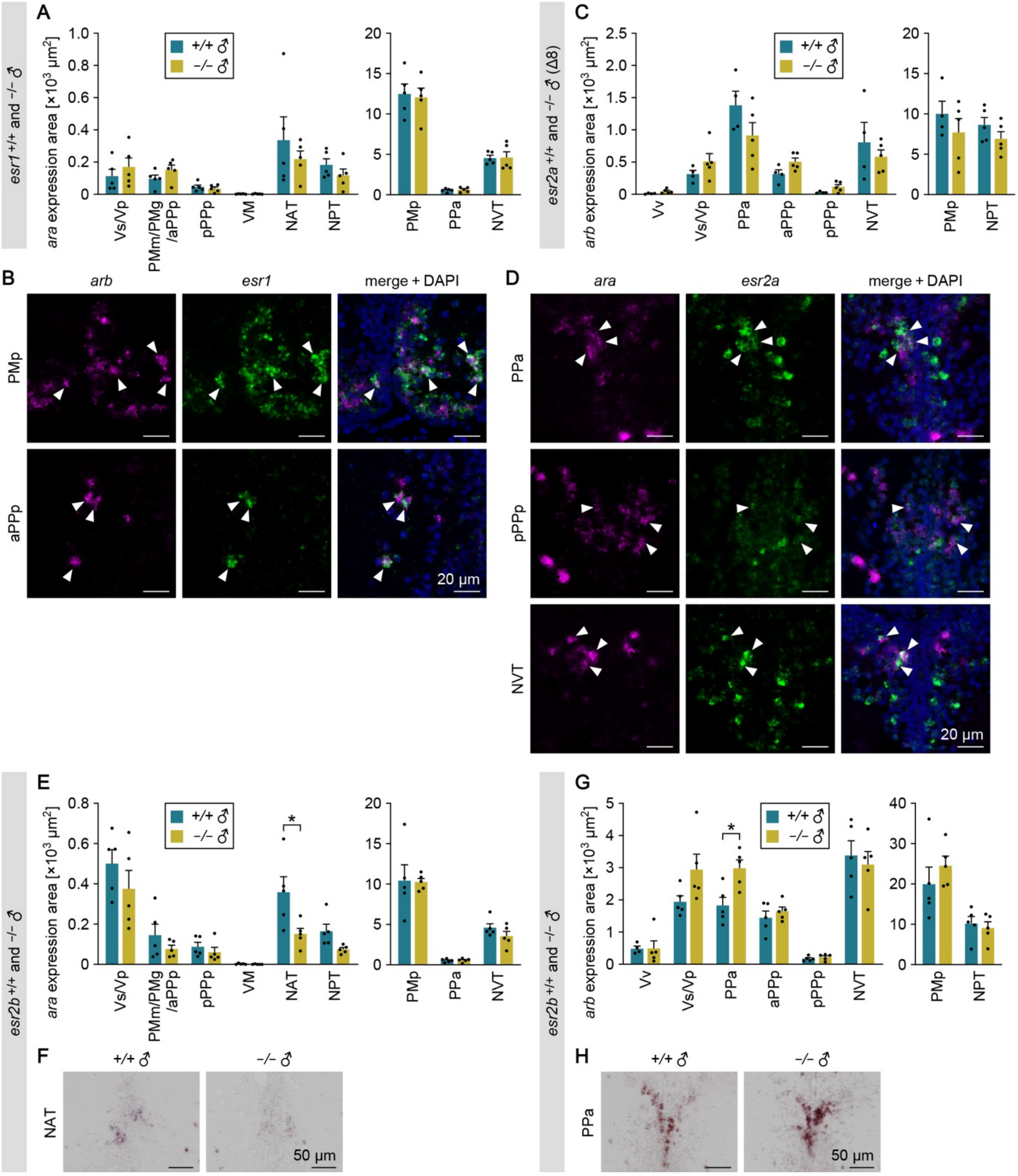


**Figure 5—figure supplement 2. Expression of *ara* and *arb* in the brain of males deficient for each ESR.**

(**A**) Total area of *ara* expression signal in each brain nucleus of *esr1*$^{+/+}$ and *esr1*$^{-/-}$ males (n = 5 per genotype). (**B**) Representative images showing the coexpression of *arb* and *esr1* in the PMp and aPPp. Left panels show *arb* expression (magenta), middle panels show *esr1* expression (green), and right panels show the merged images with DAPI staining (blue). White arrowheads indicate neurons coexpressing *arb* and *esr1*. (**C**) Total area of *arb* expression signal in each brain nucleus of *esr2a*$^{+/+}$ and *esr2a*$^{-/-}$ males (Δ8 line; n = 4 and 5, respectively, except for NPT of *esr2a*$^{+/+}$ males, where n = 5). (**D**)

Representative images showing the coexpression of *ara* and *esr2a* in the PPa, pPPp, and NVT. Left panels show *ara* expression (magenta), middle panels show *esr2a* expression (green), and right panels show the merged images with DAPI staining (blue). White arrowheads indicate neurons coexpressing *ara* and *esr2a*. (**E**) Total area of *ara* expression signal in each brain nucleus of $esr2b^{+/+}$ and $esr2b^{-/-}$ males (n = 5 per genotype except for NPT of $esr2b^{+/+}$ males and Vs/Vp of $esr2b^{-/-}$ males, where n = 4). (**F**) Representative images of *ara* expression in the NAT. (**G**) Total area of *arb* expression signal in each brain nucleus of $esr2b^{+/+}$ and $esr2b^{-/-}$ males (n = 5 per genotype except for Vv of $esr2b^{+/+}$ males, where n = 4). (**H**) Representative images of *arb* expression in the PPa. Each quantitative data set is displayed in two graphs for visual clarity. Scale bars represent 20 μm (B, D) and 50 μm (F, H). For abbreviations of brain nuclei, see Supplementary file 1. Statistical differences were assessed by unpaired *t* test, with Welch's correction where appropriate (A, C, E, G). Error bars represent SEM. $*P < 0.05$.

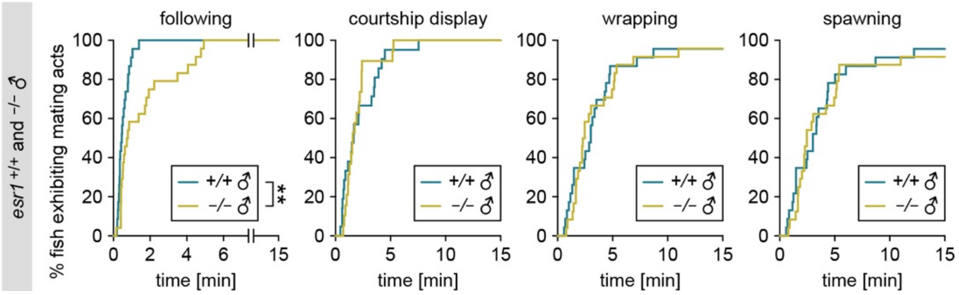


**Figure 5—figure supplement 3. Mating behavior of *esr1*-deficient males.**

Latency of *esr1*$^{+/+}$ and *esr1*$^{-/-}$ males (n = 23 and 24, respectively) to initiate each mating act toward the stimulus wild-type female. Statistical differences were assessed by Gehan–Breslow–Wilcoxon test. $^{**}P < 0.01$.

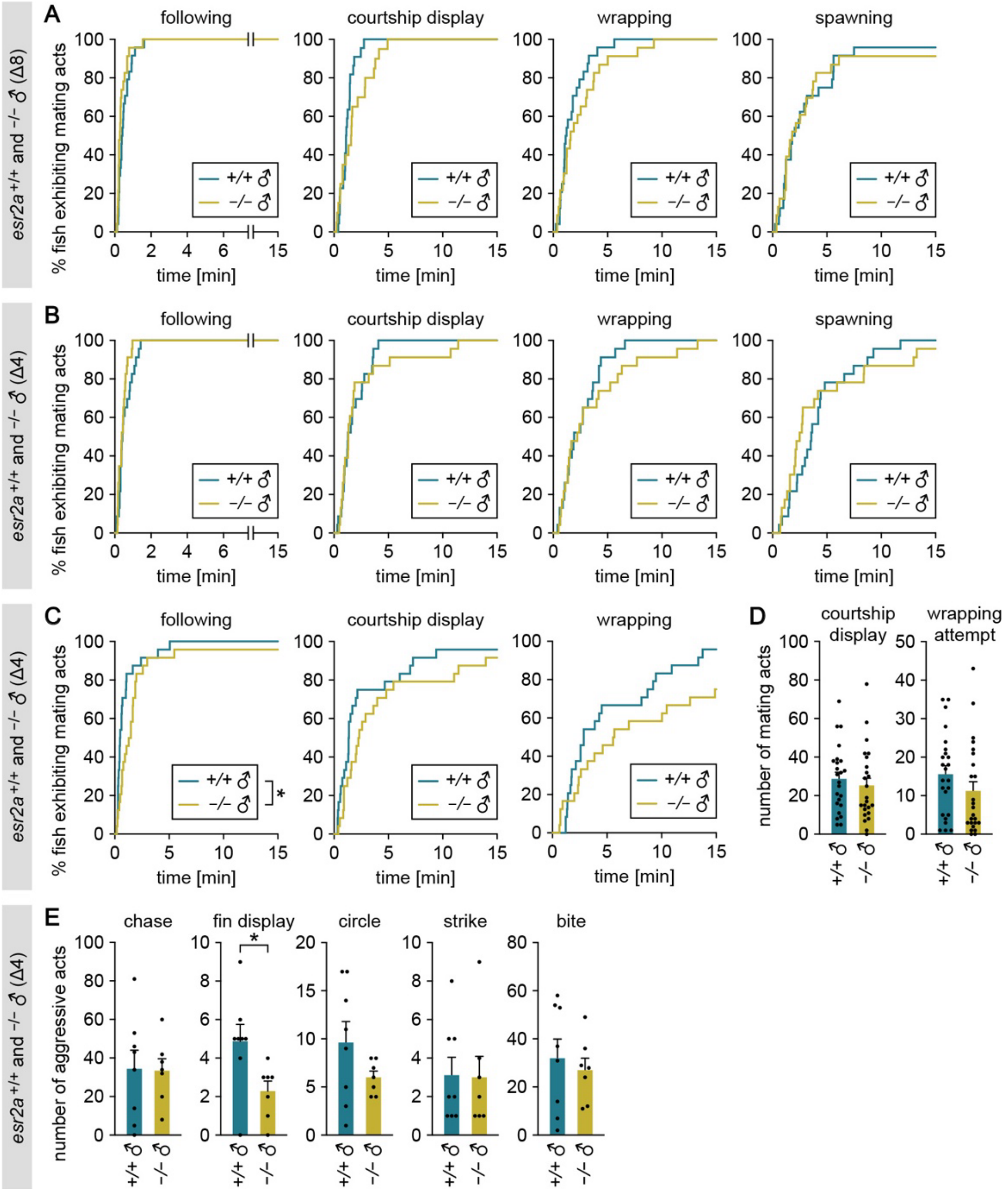


**Figure 5—figure supplement 4. Mating and aggressive behaviors of *esr2a*-deficient males.**

(**A**) Latency of *esr2a*$^{+/+}$ and *esr2a*$^{-/-}$ males (Δ8 line; n = 24 and 23, respectively) to initiate each mating act toward the stimulus wild-type female. (**B**) Latency of *esr2a*$^{+/+}$ and *esr2a*$^{-/-}$ males (Δ4 line; n = 23 per genotype) to initiate each mating act toward the stimulus wild-type female. (**C**) Latency of *esr2a*$^{+/+}$ and *esr2a*$^{-/-}$ males (Δ4 line; n = 24 per genotype) to initiate each mating act toward the stimulus *esr2b*-deficient female. (**D**) Number of each mating act performed. (**E**) Total number of each aggressive act observed among *esr2a*$^{+/+}$ or *esr2a*$^{-/-}$ males (Δ4 line; n = 8 and 7, respectively) in the tank. Statistical differences were assessed by Gehan–Breslow–Wilcoxon test (A, B, C) and unpaired *t* test, with Welch's correction where appropriate (D, E). Error bars represent SEM. *$P < 0.05$.
